# Formal cooperation in macaque monkeys obtained via single-payoff change from formal coordination games

**DOI:** 10.1101/2022.12.25.521899

**Authors:** Charlotte van Coeverden, Wolfram Schultz

## Abstract

Coordination and cooperation are hallmarks of the behavior of social animals. Coordination requires common choices to obtain maximum benefit, whereas cooperation requires to forgo immediate selfish outcome for later common maximum benefit. A well validated economic game for investigating cooperation is Prisoner Dilemma (PD). Recent studies show that monkeys cooperate to a limited extent when playing an iterated PD. In our experiment, macaque monkeys made choices on a touchscreen to obtain juice reward whose amount depended on the choices of both animals. We designed four coordination games and two cooperation games (iterated PD) that differed only in a single payoff (the so-called temptation) while all other payoffs remained constant. The increasing temptation payoff resulted in performance that varied somewhat in the coordination game (probability of common choice between p = 0.55 and p = 0.70) but dropped in both cooperation games while nevertheless remaining significant (p = 0.28 to p = 0.68). The response time of the second player increased significantly when the first player chose the cooperative option across all games, suggesting reciprocation; further, the animals seemed to benefit from seeing the action of the other player, indicating that the choices incorporated a social component. Taken together, our results demonstrate good cooperation in the iterated PD by macaque monkeys after being primed with coordination games.

## Introduction

Cooperation characterizes social behavior that results in the common good and thus is of societal importance. Cooperation has two key characteristics: players make common choices that allow them to obtain the ultimate highest collective gain, but they need to forego an immediate higher selfish gain. Thus, cooperation results in an ultimate optimal collective benefit by outweighing an initial suboptimal individual gain (Nowak 2006). By contrast, coordination characterizes common choices for maximal gain without foregoing an immediate selfish gain. Thus, coordination results in immediate optimal collective outcome without foregoing an initial selfish gain. Overall, by acting beyond the short term, cooperation is more demanding than coordination.

The quintessential abstraction and operational definition of cooperation is provided by the Prisoner’s Dilemma (PD) that had originally been designed for performance in single trials. The most beneficial choice for an individual player in such single-shot PD is selfish defection in which the other player receives only a small payoff, thus resulting in limited common benefit. However, social behavior usually depends on experience and thus involves repeated choices during which long-term gains can be evaluated. Correspondingly, monkeys in the laboratory make repeated choices, in particular in tasks suitable for neurophysiology with appropriate statistics. The repeated experience is captured in the iterated PD in which players perform an unknown number of PD repetitions. The players come to cooperate over repeated trials, as the long-term common outcome exceeds the short-term individual gain, and the short term losses from getting defected by a selfish player have a chance to give way to an overall mutual gain for both players. All players in iterated and unlimited PD gain over the long run by cooperating. Thus, iterated PD captures well the notion of cooperation: two players obtain the highest common outcome by foregoing higher short-term individual gains (Rapoport 1974). PD-style cooperation games contrast with coordination games in which each of two players chooses the best option without having to forego any initial higher gain.

Cooperation and coordination are not unique to humans and have been studied in animals (Stephens et al. 2002). In iterated versions of these games, macaque and cebus monkeys perform well in coordination games (de Waal & Davis 2003; Brosnan et al. 2012; Smith et al. 2019) but are just above chance level or instable in cooperation games (iterated PD; Haroush & Williams 2015; Smith et al. 2019). The results indicate that macaque monkeys can coordinate their choices for a common good but have difficulties cooperating in formal iterated PD. As birds and rats do not usually cooperate in straightforward iterated PD (Gardner et al. 1984; Clements & Stephens 1995; Stevens & Stephens 2004), cooperation may evolve across species. Thus, the evolutionary aspect and the societal importance of cooperation warrants further exploration of the capacity of cooperation in monkeys.

Our study asked how macaque monkeys can come to cooperate more efficiently, using the formal iterated PD in which reward depends on the combined choices of two animals. We benefitted from a previous approach that achieved cooperation in birds by priming them with maximum reward for common choice (Stephens et al. 2002). Our monkeys played coordination games in which the benefit from choosing the ‘Cooperate’ option and the disadvantage from choosing the ‘Defect’ option were particularly well visible. We tested a series of coordination games that differed only in a single payoff (the so-called ‘temptation’ payoff for choosing ‘Defect’ instead of ‘Cooperate’). We stepwise increased the temptation payoff across games until the coordination games became cooperation games (iterated PD). The temptation payoff encourages immediate defection and thus discourages cooperation, which challenges an agent’s appreciation of long-term gain from cooperation. Thus, we translated the easier ability of immediate coordination into the more demanding ability of cooperation for common long-term gain. We tested these games in monkeys, as they are the species of choice for ultimate investigations of neuronal mechanisms of sophisticated social behavior. The animals made cooperative choices with probabilities between p = 0.28 and p = 0.68 that significantly exceeded chance (p = 0.25). These results should be useful for investigating neuronal mechanisms of social behavior beyond the already known coding of other’s actions and choices (di Pellegrino et al. 1992; Hosokawa & Watanabe 2012; Báez-Mendoza et al. 2013; Haroush & Williams 2015; Chang et al. 2015; Grabenhorst et al. 2019; Báez-Mendoza et al. 2021).

## Methods

### Animals and ethics

Three adult male rhesus monkeys (Macaca mulatta) served in this study. Monkey L40 (‘Monkey L’), weighing 12.0 – 13.6 kg, was trained beginning May 25, 2014 and tested in the full task from Sep 8, 2014 to May 22, 2017. Trident (‘Monkey T’), weighing 8.9 – 14.2 kg, was trained beginning May 25, 2014 and tested in the full task from Sep 8, 2014 to Feb 8, 2017. Virtue (‘Monkey V’), weighing 5.1 kg – 9.0 kg, was trained beginning August 14, 2015 and tested in the full task from Oct 28, 2015 to May 22, 2017. Thus, all three animals were tested in the full task simultaneously from August 14, 2015 to Feb 8, 2017.

This research has been ethically reviewed, approved, regulated and supervised by the following individuals and institutions in the UK and at the University of Cambridge (UCam): the Minister of State at the UK Home Office, the Animals in Science Regulation Unit (ASRU) of the UK Home Office implementing the Animals (Scientific Procedures) Act 1986 with Amendment Regulations 2012, the UK Animals in Science Committee (ASC), the UK Home Office Inspector for our UCam laboratory, the UK National Centre for Replacement, Refinement and Reduction of Animal Experiments (NC3Rs), the UCam Animal Welfare and Ethical Review Body (AWERB), the UCam Governance and Strategy Committee, the Home Office Establishment License Holder of the UCam Biomedical Service (UBS), the UBS Director for Governance and Welfare, the UBS Named Information and Compliance Support Officer, the UBS Named Veterinary Surgeon (NVS), and the UBS Named Animal Care and Welfare Officer (NACWO).

The animals were housed together with other adult male rhesus monkeys in home cages with enrichment, as follows: Monkeys T and V were cage mates together with two other monkeys; Monkey L was housed in a different holding room in a cage of his own together with two to three other monkeys in the same room. Before behavioral testing, Monkeys T and L had a headpost that had been surgically implanted for later neuronal recordings using general anesthesia and aseptic procedures (the early implantation served for allowing gradual osteointegration); however, the currently described behavioral tests were performed without head fixation. When seated in their respective primate chairs, Monkey L dominated Monkey T who dominated Monkey V, as tested by their tendency to grab food morsels offered by an experimenter; the dominance hierarchy appeared to correlate with their respective body weights. In the common home cage of Monkeys T and V, another monkey not used for the current experiments dominated Monkey T (who dominated Monkey V).

For each daily session, the animal entered from their home cage into an individually adjusted and comfortably fitting purpose-made primate chair (Crist Instruments); each animal had its own primate chair for the duration of the experiment. The chair with the animal inside was wheeled into the laboratory, where the animal was trained in three pair-wise combinations (dyads) with one of the other experimental monkeys to perform the behavioral task in which it received juice reward by contacting a touch key and subsequently touching a horizontally mounted touchscreen. After an initial habituation of several months to the primate chair, laboratory, touch key, touchscreen and juice spout, the animals were trained in the specific social task.

### Behavioral task

Two monkeys in any of the three possible dyads faced each other across the common horizontal touchscreen. Both animals performed in a binary choice task for juice reward (Fig. 1A). A trial started when both animals contacted a touch-sensitive key, upon which the same two fractal stimuli appeared on the touchscreen for each animal. The two fractals indicated the two choice options; they had grey color, equal size, normalized luminosity and were visible to both animals (unless otherwise stated). At 500 ms after fractal presentation, a small grey rectangle appeared next to each fractal to elicit each animal’s choice. The animal released the touch key and selected the target of its choice. Each animal was free to choose at a self-determined moment but had to complete the trial within four seconds. Thus, there was neither a specific signal nor any other requirement that determined the sequence in which the animals chose. Although the limited response time encouraged rapid responses, the duration was long enough to allow an animal to observe the other animal’s choice before making its own choice. When both animals had chosen within the four-second time window, they received specific numbers of 0.15 ml drops of blackcurrant juice depending on each choice option. A computer-controlled solenoid valve (SCB262C068; ASCO) delivered the juice at a 150 ms interval between the two animals from a spout in front of each animal’s mouth.

**Fig. 1.**
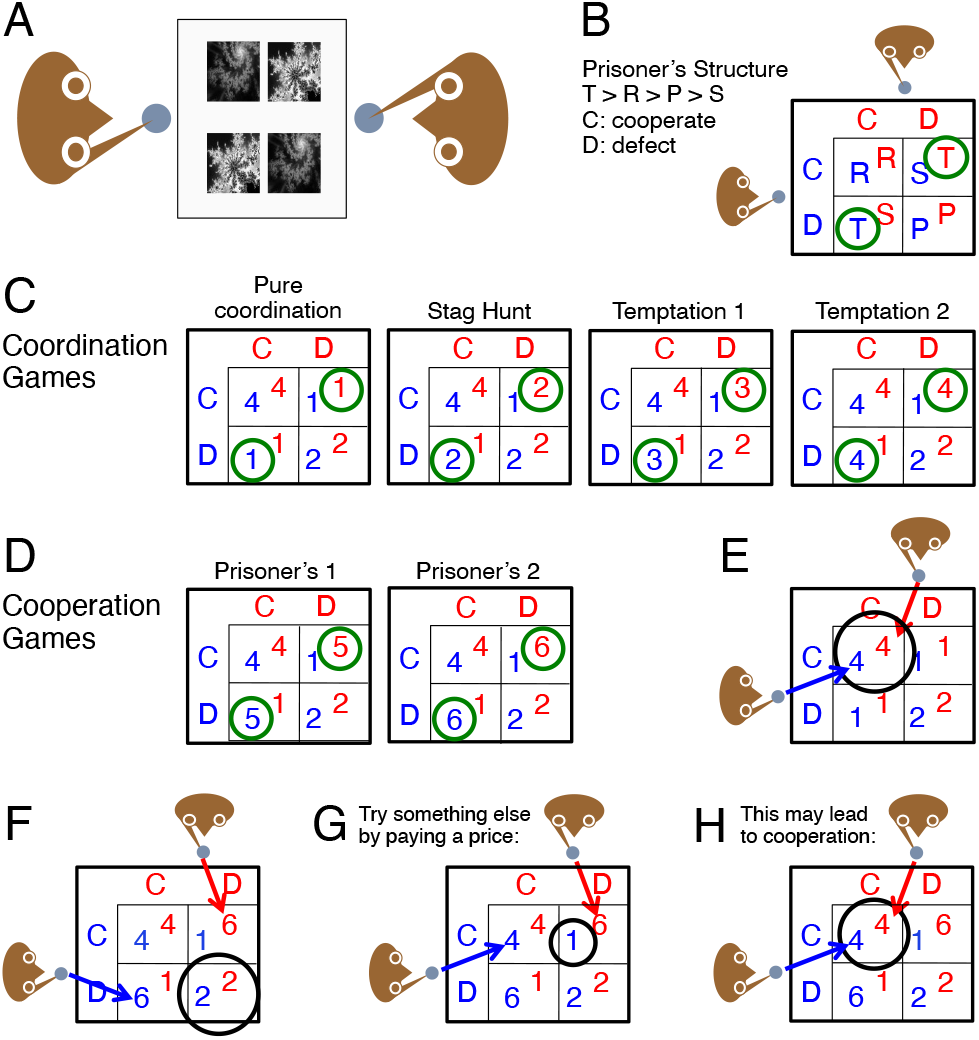
Behavioral task and experimental design. Change of a single payoff mediates the transition from formal coordination games to formal cooperation games (Prisoner’s Dilemma, PD). (*A*) Task setup with two monkeys facing each other across a horizontally mounted touchscreen. Each animal lifted one hand off a touch sensitive holding key to touch one of two grey fractals indicating different amounts of reward juice. (*B*) Payoff matrix defining the cooperation game. Payoffs are ranked as T (temptation) > R (reward) > P (punishment) > S (getting suckered). D: defection; C: cooperation. (*C*) Payoff matrices of coordination games. Green circles indicate the single payoff that distinguishes each game and varies across games (T, temptation). The second game from left is also known as Stag Hunt. (D) Payoff matrices of formal cooperation games. The coordination game becomes a coordination game when the temptation payoff (T) surpasses the reward payoff (T > R; green circles), making defection (D) more valuable than cooperation (C). (*E*) Playing a coordination game. The players obtain maximal payoff by both choosing the individually best option (coordination; circle). (*F*) Playing an iterated PD cooperation game. Each player is tempted to choose the largest payoff (6 units). However, when both players do so, each player receives only a low payoff (2 units; circle). (*G*) Unsatisfied, in the same cooperation game, one player may instead choose the other option that would have provided a smaller payoff had both players chosen it (4 units) but receives only 1 unit because of the other player’s choice (1 unit; circle). (*H*) In the same cooperation game, if the other player also changes the choice, both players obtain the maximum common good (4 units for each player; circle).

To facilitate consideration of each option and prevent overtraining, a new set of two fractals was introduced after a given fractal set had been used for several trials (284 ± 185 trials per fractal set; mean ± standard deviation). Each fractal set was used only for one specific game. Thus, a new fractal set indicated either continuation of the same game or use of a new game. This procedure aimed to ensure exploration of both options before committing to a single option. All games were presented in pseudorandom order.

### Games

The properties of economic games are determined by each game’s specific payoff matrix. The payoff from choosing a particular option depends on the own choice and, importantly, also on the other player’s choice. The PD game is special in paying the highest payoff for a deliberately different choice than an opponent who is choosing an option that would lead to maximal common payoff when both players were to choose the same option (‘cooperation’). By contrast, in a coordination game, both players receive the highest payoff when they both choose the best option and, importantly, cannot get more payoff by choosing differently. All our games contained two choice options that were identical for both animals and were labelled option C and option D (alluding to ‘cooperate’ and ‘defect’ with PD). To simplify the task for the monkeys, we used only positive outcomes, namely drops of rewarding fruit juice.

The characteristics of the PD game are as follows. When both players choose to cooperate (C), they receive an intermediate payoff (R for reward). When both players choose to defect (D), they receive a smaller intermediate payoff (P for punishment). When only one player chooses defect (D) and the other player chooses C (cooperate), the defecting player receives the highest payoff (T for temptation) and the cooperating player receives the lowest payoff (S for getting suckered). Thus, the hierarchy of payoffs in PD is defined as (T = temptation) > R (= reward) > P (= punishment) > S (= getting suckered) (Fig. 1B).

When playing a coordination game, both players receive the highest payoff when they both choose the cooperation option C (for simplicity called ‘cooperation’ even for coordination games) and the same or lower payoff when both choose the defection option D. When each player chooses a different option, they receive a submaximal payoff that is smaller than when both players choose option C.

We systematically advanced the animals’ training and testing from four coordination games to two PD games by stepwise incrementing a single payoff specific for each game (Fig. 1C and D, green circles). The defining payoff was labelled as T (temptation) in correspondence to PD. It was paid out to the player choosing option D when the other player chooses option C. The temptation payoff T defined the difference between coordination games and cooperation games: payoff T in coordination games was never higher than the payoff received when both players chose the same option, whereas payoff T in cooperation games was the highest payoff and exceeded the payoff for any same choice. Thus, the stepwise increase of the defining temptation payoff T defined the transition from coordination games (Fig. 1C) to cooperation games when T became the highest payoff (Fig. 1D). As the word indicates, the higher payoff for defecting on a cooperating player constitutes a temptation that defines the PD. Thus, we were able to transition from coordination games to cooperation (PD) games by varying a single payoff while holding all other payoffs constant.

### Analysis of choices

We recorded choices from the animals in all three dyads. We tested whether choices by one animal were more likely based on the already known choices of the other animal, using trials in which the animal had chosen second. We used the following logistic regression:

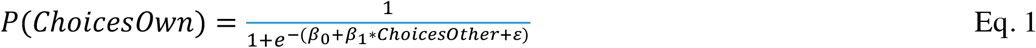

### Analysis of strategies

Strategies can be summarised as sets of rules that determine the choice given a set of circumstances. Trials were classed according to three strategies for each animal: Perseverance (STAY), Win-stay Lose-Shift (WSLS) and Tit-for-Tat (TFT). All strategies depended on combinations of current and previous trial (CT and PT) of own and other’s choices and payoffs. We used the following definitions:

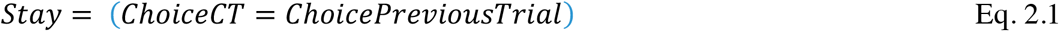

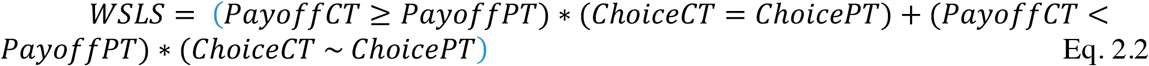

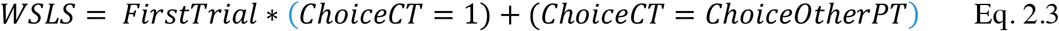

The effect of strategies on cooperation was analysed by means of logistic regression. As the strategies were too highly correlated to use them in one model, their effect on cooperation was assessed separately using different models.

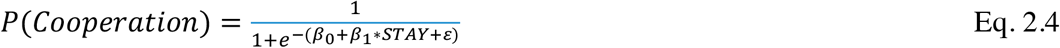

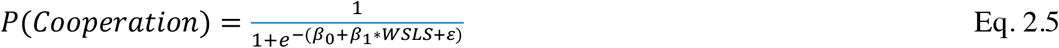

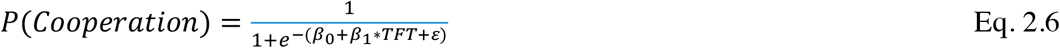

Statistics were calculated with SPSS (IBM) and Excel (Microsoft). Matlab (Mathworks) and SPSS (IBM) were used for all regression analyses.

## Results

### Patterns of coordination and cooperation

Using a total of three monkeys, we studied choices in three pairs of two monkeys each (dyads) that sat opposite to each other across a horizontally mounted computer touchscreen. The two animals chose near-simultaneously, without prescribed inter-individual sequence, between two simultaneously presented options (C for cooperate, D for defect); their payoffs were indicated by grey-scale fractal stimuli (Fig. 1A, B) (note that option C denotes the coordination option in coordination games but the cooperation option in cooperation games; for simplicity, we call C commonly ‘cooperation’ in this study). We inferred the animal’s stochastic preference from the probability of choosing one option over all other options of the same option set (Luce 1959). However, in these social games, the rewards for each animal depended on both the own and the other’s choice, and the reward was not fully predictable from the choice of a particular player alone.

The three monkeys were trained in six formal games in chronological sequence (although the games were tested in pseudorandom alternation after full training). We changed the single payoff for the temptation payoff (T) in steps of one unit, thus transitioning from coordination games (Fig. 1C, green circles) to cooperation games (Fig. 1D). With our settings, the formal coordination game became a formal cooperation game when the temptation payoff exceeded the reward payoff (T > R), which implemented the temptation that challenges the coordinated action as essence of cooperation. According to general assumptions, the propensity of both players to choose the cooperation option (C) increases with higher reward payoff (R) and lower temptation payoff (T) (Camerer 2002).

The difference in performing in these two classes of games is illustrated in Fig. 1E-H. In a coordination game, reward is maximized when both players choose simply the best-paying option for each player (Fig. 1E). This strategy contrasts with the choices in a typical iterated PD cooperation game. When both players attempt to choose the largest payoff for themselves, they receive only a suboptimal total payoff (Fig. 1F; circle: 2 units each). Alternatively, when the first player chooses the largest payoff (6 units, red player in Fig. 1G), the other player may choose the other, smaller payoff but receives only a very small payoff (circle: 1 unit); the total payoff for the two players is still suboptimal (6 + 1 units). But the second player’s choice of the smaller payoff may prime the first player to also try that smaller payoff, which then leads to the highest total payoff (Fig. 1H; circle: 4 + 4 units). Thus, instead of one player gaining high and the other player gaining low, both players obtain the highest total payoff by both choosing an option with submaximal individual payoff.

The three monkeys performed a total of 40,903 dyad trials. In all four coordination games (T, temptation payoffs 1-4), the three animals learned rapidly to coordinate their actions and consistently chose together the more rewarded option C (Fig. 2A). By contrast, the choices varied considerably in the PD game (Fig. 2B). For example, with T set to payoff 5, Monkey L consistently failed to cooperate and defected on Monkey T across a whole daily session while Monkey T was lenient and stubbornly offered to cooperate (Fig. 2B, leftmost graph). In another session, Monkey L cooperated initially but came to defect as Monkey T defected almost consistently and only at session end started to cooperate (next graph); apparently Monkey L ran out of patience with the defecting Monkey T. When playing with Monkey T at temptation payoff T = 5, Monkey L cooperated consistently even though Monkey T defected initially (next graph); thus, the tolerant behavior of Monkey L may have led Monkey T to cooperate. In another session, both monkeys cooperated well with only short bouts of defection (rightmost graph). Further graphs confirm the rapid and consistent development of coordination (Fig. 2C) but the slower and more variable developing cooperation (Fig. 2D). Thus, the initial variable performance in the coordination games suggests that the animals explored the contingencies for a short period. The initial period was followed by a longer sustained period of stable cooperative choices. Nevertheless, compared to performance in coordination games, cooperative games involved more switching between choice options and larger chances of sustained defection, especially at first or early sessions.

**Fig. 2.**
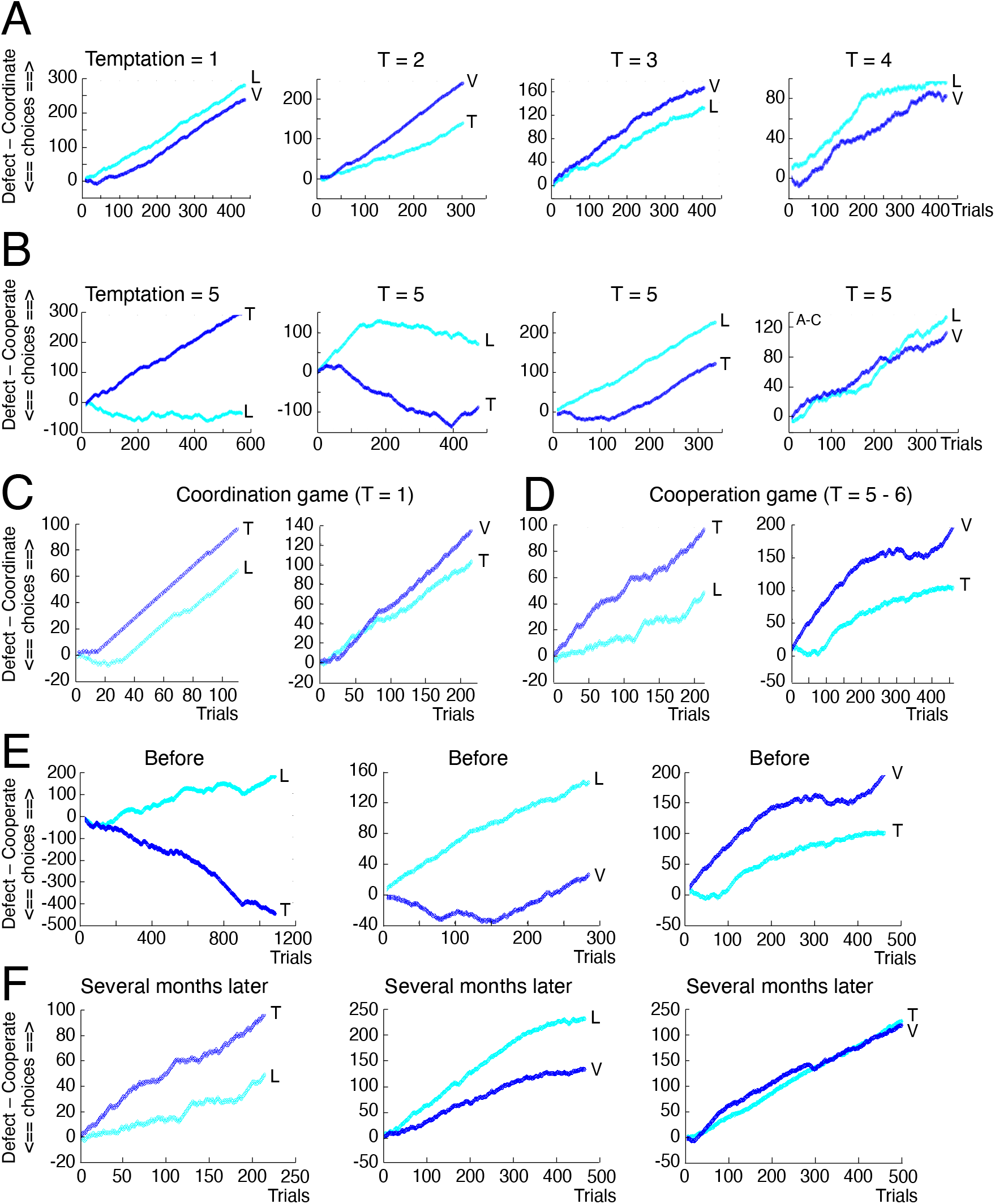
Patterns of coordination and cooperation. (*A*) Good performance in all coordination games tested across successive trials within single sessions. After very brief exploration, the animals chose the optimal outcome (mutual coordination). The payoff setting of option T (temptation to defect) defined the specific coordination game, whereas all other payoffs were identical. In a coordination game, the payoff T for asymmetrical choices between animals is never larger than the payoff R for same choices (reward R for coordination) between the two choosers (payoffs T ≤ R). L, T and V refer to the three monkeys tested; L / V and V / T refer to two dyads. (*B*) Challenging cooperation in Prisoner’s Dilemma (PD). PD has higher payoff for temptation T compared to reward R (T > R), making defection beneficial for the defector. Graphs show development of cooperation (increasing slope), alternation between cooperation and defection (approximately horizontal slopes) and defection (decreasing slope) across session trials, with payoff T of 5. Thus, the behavior can be described, from left to right, as leniency and consistent offer to cooperate in the face of the other animal’s defection, initial cooperation turning into defection as the other animal defected, unilateral offer to cooperate leading to mutual cooperation, and good mutual cooperation. (*C*) Immediate and reliable coordination in two other dyads (T / L and V / T). (*D*) Slowly developing cooperation in PD. (*E - F*) Development of cooperation in PD after several months of experience (T = 6) in all three dyads (L / T, L / V and T / V). The initial hopeless or variable cooperation (*E*) turns ultimately into more consistent cooperation several months later in the same dyads (*F*).

Given the variability of performance in the cooperation (PD) games (T = 5 or T = 6), we investigated whether extended experience might lead to better cooperation. All three dyads showed initial difficulties in cooperation, with persistent defection even in the face of cooperation by the opponent monkey (Fig. 2E left), considerable initial hesitation (Fig. 2E center), or only moderate cooperation (Fig. 2E right). However, cooperation increased significantly with all three dyads over 3-5 months of testing on several days each week (Fig. 2F; from 10% to 50% of trials; P = 0.09 E-30), whereas coordination was high early on and did not increase further (50 – 60% of trials; P = 0.086). Thus, cooperation developed more slowly than coordination but ultimately succeeded.

### Development of cooperation

The data presented so far demonstrate that monkeys can cooperate in PD but that cooperation takes time to develop (Fig. 2E, F). As one of the aims of the current experiments was to investigate more explicit ways to foster cooperation, we designed our gambles such that a single payoff (T for temptation) defined each specific game and that the stepwise increment of that payoff changed coordination games into cooperation (iterated PD) games (Fig. 1C, D). In all four coordination games, the animals showed good levels of commonly choosing the coordination (C) option that maximized their reward. In Fig. 3A, the good coordination was evidenced by high probabilities of each animal of each dyad consistently choosing the coordination (C) option (close to top right corners in top four heatmap rows). When the payoff change for temptation transformed the coordination games into cooperation games (from T = 1 - 4 to T = 5 and 6), both animals chose the cooperation option C less consistently, as indicated by more widely scattered choice probabilities (Fig. 3A, two bottom rows).

**Fig. 3.**
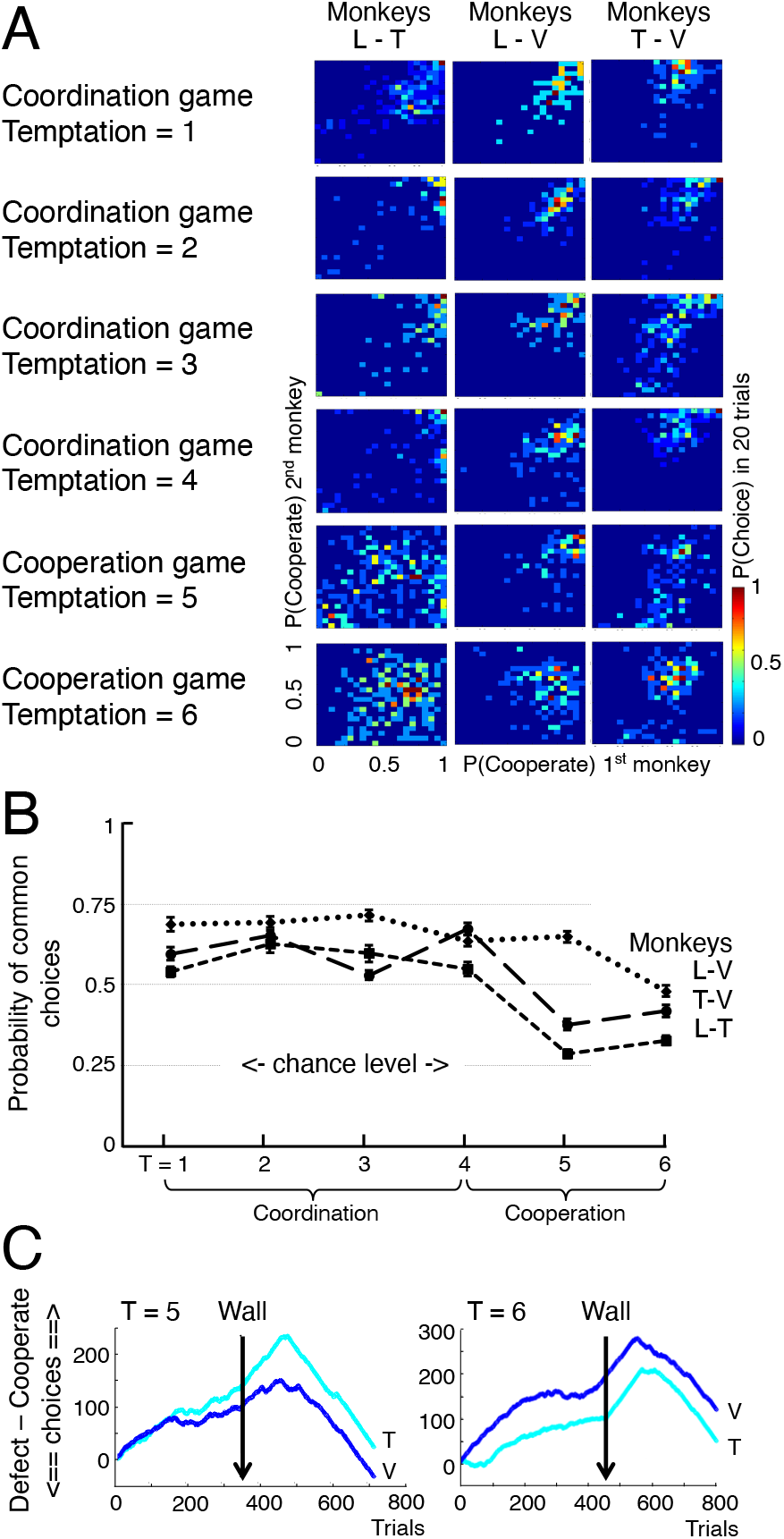
Experience with coordination leads to emergence of cooperation. (*A*) Heatmaps of choice probabilities for each monkey of all three tested dyads. The single temptation payoff (T) defined the difference between the four coordination games and two cooperation games. Rainbow colored dots concentrated at the top right corner of each plot indicate good coordination and cooperation between the animals. (*B*) Summary of coordination and cooperation levels, as indicated by the probability of choosing the cooperation (C) option over the defection (D) option. In all dyads, performance was consistently high in all coordination games but decreased somewhat in cooperation games in which choices still differed significantly from chance. T, temptation payoff. (*C*) Wall control. Breakdown of cooperation when a wall between the two monkeys T and V prevented visual inspection of each other’s options and choice (while maintaining other visual contact between the animals).

Different quantification of the same data demonstrated high coordination levels in the four coordination games (T = 1 – 4; probability of both animals choosing coordination between p = 0.55 and p = 0.70) that decreased somewhat to still significant, consistent and above-chance levels in both cooperation games (T = 5 and 6; probability of both animals choosing cooperation between p = 0.28 and p = 0.68) in all three dyads (Fig. 3B) (p < 0.01). The dyad L – V showed lower cooperation with the temptation payoff increasing from T = 4 to T = 5, which confirmed the notion that cooperation decreases with increasing temptation payoff (T) (Camerer 2002), whereas the other two dyads failed to show substantial changes. These data suggest overall excellent performance in the coordination games, and slightly less but nevertheless significant cooperation in cooperation games (iterated PD) despite the well-rewarded temptation to defect.

### Importance of viewing each other’s options and choice

To assess the influence of viewing each others choices, we temporarily placed a 12 cm high wall in the center of the horizontal touchscreen between the animals. The wall obstructed the view of the touchscreen and thus made it impossible for each animal to view the other animals’ options and choices. The animals were still able to see each other’s hand on the touch key (but not the touchscreen) and the delivery of the reward after the choice, as well as task-unrelated arm and face movements. The view of the remainder of the laboratory, the behavioral setup, the behavioral task, the fractal stimuli predicting the payoffs, and the payoffs themselves remained unchanged. Thus, the animal could only infer the other animal’s choice from seeing the reward the other animal received while taking into account his own choice.

Despite good performance in the cooperation game with both temptation values before wall placement, cooperation broke down within about 100 trials after wall placement and was largely replaced by defection in both animals of the tested dyad (Fig. 3C). Thus, viewing each other’s options and choice seemed to be crucial for mutual cooperation.

### Offer and reciprocation

The monkeys were free to choose as soon as the two fractals for each animal appeared. The unconstrained order of choice allowed separate analysis according to first or second chooser. When Monkey L chose first in the coordination game, he chose the cooperation option C significantly more frequently than the defection option D; the same was true for Monkey T (Fig. 4A, blue to yellow). The tendency declined and became variable in the cooperation game (orange and red). Only Monkey L chose cooperation more frequently than defection and thus offered above-chance cooperation despite facing potential defection by the second chooser. By contrast, Monkey T mostly chose defection as first mover.

**Fig. 4.**
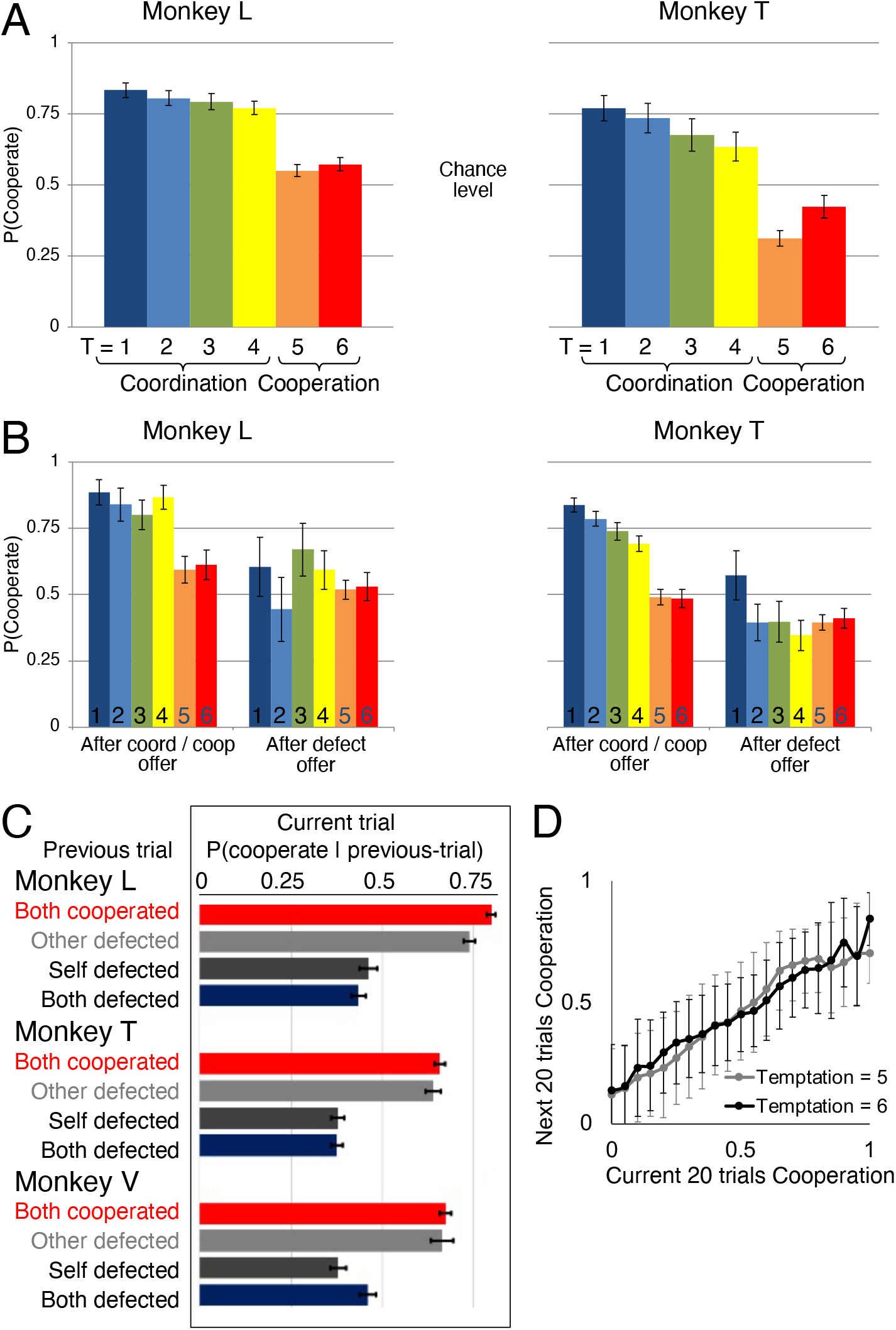
Offer and reciprocation. Monkeys were free to choose after the simultaneous appearance of the two options, which allowed post-hoc distinction between first and second chooser. (*A*) Probability of selecting the cooperation option by first chooser. Error bars are 95% confidence intervals. T, temptation payoff. (*B*) Probability of selecting the cooperation option by second chooser after first chooser had selected cooperation (left) or defection (right) in same trial. (*C*) Choice of cooperation option in previous trial lead to more cooperation in current trial (temptation payoff T = 5 or T = 6). (*D*) Current cooperation induced future cooperation.

When analyzing the choices of the second-choosing animal, we found that both Monkeys L and T chose the cooperation option C in the four coordination games significantly more frequently than the defection option D when the first mover had also chosen cooperation (Fig. 4B, blue to yellow). The proportion of choices of coordination dropped significantly when the first mover had chosen defection (odds ratios for monkey L: 2.1, 6.8, 3.8 and 4.3 for temptation payoffs 1-4, respectively, all Wald Statistics significant with p < 0.01; odd ratios for monkey T: 3.4, 3.5, 2.6 and 3.4 for temptation payoffs 1-4, respectively, all Wald statistics significant with p < 0.01). A similar reduction was seen with the overall lower cooperation in the cooperation game. Cooperation by the second mover was significantly higher after the first mover had chosen the cooperation option, as compared with the first mover having chosen the defection option (orange and red) (odds ratios for monkey L: 1.6 and 1.5 for temptation payoffs 5 and 6, respectively, both Wald Statistics significant with p < 0.01; odd ratios for monkey T: 1.4 for both temptation payoffs 5 and 6, both Wald statistics significant with p < 0.05). Thus, the choice of the first-choosing monkey affected the other monkey’s choice in all games. In particular, cooperation seemed infectious with both animals.

Cooperation in PD depended in all three animals also on experience in the previous trials. Cooperation of both animals in the previous trial resulted in high probability of choosing the cooperation option C again (Fig. 4C, red). By contrast, choice of the cooperation option was less likely in trials following any defection (grey and black). Correspondingly, the degree of mutual cooperation in the current 20 trials correlated with the degree of mutual cooperation in the next 20 trials (Fig. 4D) (Monkey L: beta = 0.57, F(6,994,1) = 3484.8, p < 0.001; Monkey T: beta = 0.59, F(6,304,1) = 3,304.4, p<0.001; Monkey V: beta = 0.68, F(5,626,1) = 4,889.9, p < 0.001). Thus, previous cooperation increased current cooperation, even when own defection could result in an immediate higher payoff. It seemed that the benefits from experienced cooperation encouraged further cooperation.

### Consequences of coordination and cooperation

Reward gain increased with the increasing probability of mutual coordination and mutual cooperation in two of the three animals (Fig. 5B), both in current 20 trials (Monkey L: beta = 1.78, F(7,014,1) = 2,112.9, p < 0.001; Monkey T: beta = 1.26, F(6,324,1) = 697.35, p < 0.001; Monkey V: beta = −0.11, F(5,646,1) = 6.75, p = 0.009) and in future 20 trials (Monkey L: beta = 0.99, F(6,994,1) = 539.85, p < 0.001; Monkey T: beta= 1.09, F(6,304,1) = 503.44, p < 0.001; Monkey V: beta = −0.01, F(5,626,1) = 0.08, p = 0.8). Thus, both coordination and cooperation became more beneficial with more frequent reciprocation.

**Fig. 5.**
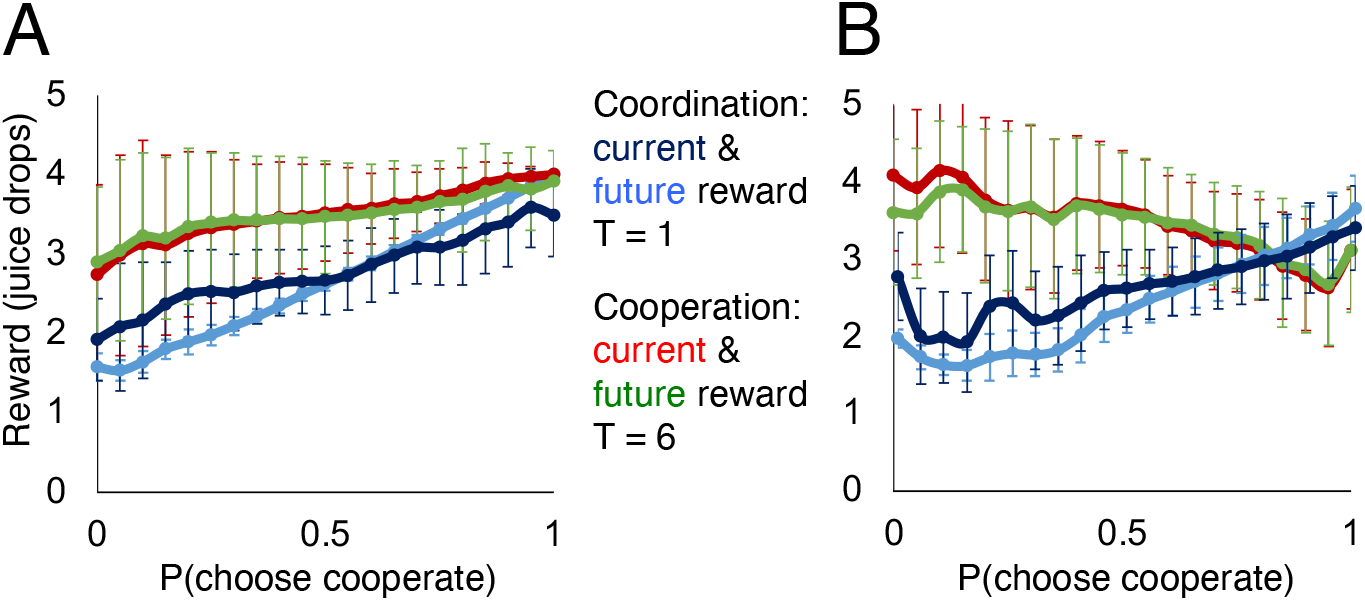
Effects of offered and mutual coordination and cooperation on current and future reward gain. (*A*) Effects of mutual choice. Higher probability of mutual coordination (T = 1) or cooperation (T = 6) increased reward gain in current and future blocks of 20 trials. (*B*) Effects of first choice. Higher probability of offering coordination by first chooser increased reward gain in current and future blocks of 20 trials (dark and light blue; temptation payoff T = 1), whereas higher probability of offering cooperation decreased reward payoff (red and green; T = 6). Thus, offering to cooperate had a price.

Closer analysis of reward gain demonstrated differential consequences between coordination and cooperation games when choosing first. Current and future reward payoff increased with the increasing probability of the first chooser offering coordination in the most simple coordination game (T = 1) (Fig. 5B, dark and light blue). The result was confirmed by regression analysis of 20 current trials and future 20 trials. Thus, offering coordination was beneficial.

In contrast to the gain by coordination, reward payoff decreased with the increasing probability of the first chooser offering cooperation in the most tempting cooperation game (temptation T = 6) (Fig. 5B, red and green). The result was confirmed by regression analysis of 20 current trials (Monkey L: beta = −1.05, F(7,014,1) = 562.8, p < 0.001; Monkey T: beta = −1.09, F(6,324,1) = 543.7, p < 0.001; Monkey V: beta = −1.33, F(5,646,1) = 1391.2, p < 0.001) and future 20 trials (Monkey L: beta = −0.96, F(6,994,1) = 457.2, p < 0.001; Monkey T: beta = −0.4973, F(6,304,1) = 105.05, p < 0.001; Monkey V: beta = −0.8933, F(5,626,1) = 562.61, p < 0.001). Thus, unconditionally offering cooperation had a price; more frequently offered cooperation may lower reward gain.

### Strategies in coordination and cooperation games

We analyzed three strategies of choosing from trial to trial: persistence of choosing the same option again (STAY), win-stay lose-shift (WSLS) and tit-for-tat (TFT) (Eqs. 2.1 – 2.6). Choices under STAY and WSLS refer primarily to own previous choice (with modulation by social choice with WSLS), whereas TFT refers only to the other’s choice and thus is outright social. Overall, the animals used WSLS more often than the other two strategies. Specifically, Monkey L consistently used STAY and WSLS more frequently than TFT, and Monkey T used mostly WSLS (Fig. 6A). The transition from coordination to cooperation games saw a moderate drop in using the non-social STAY and limited-social WSLS strategies but a sharp drop in the outright-social TFT strategy (from 63% - 74% with temptation values of 1 – 4 to 53% – 55% with temptation values of 5 and 6), although TFT performance still significantly exceeded chance. Interestingly, the use of both WSLS and TFT consistently increased cooperation more than STAY in both versions of the cooperation game (T = 5 and 6) in Monkeys L and T, with TFT being the effective strategy in Monkey T (Fig. 6B). Thus, all three animals had learned strategies that increased their cooperation.

**Fig. 6.**
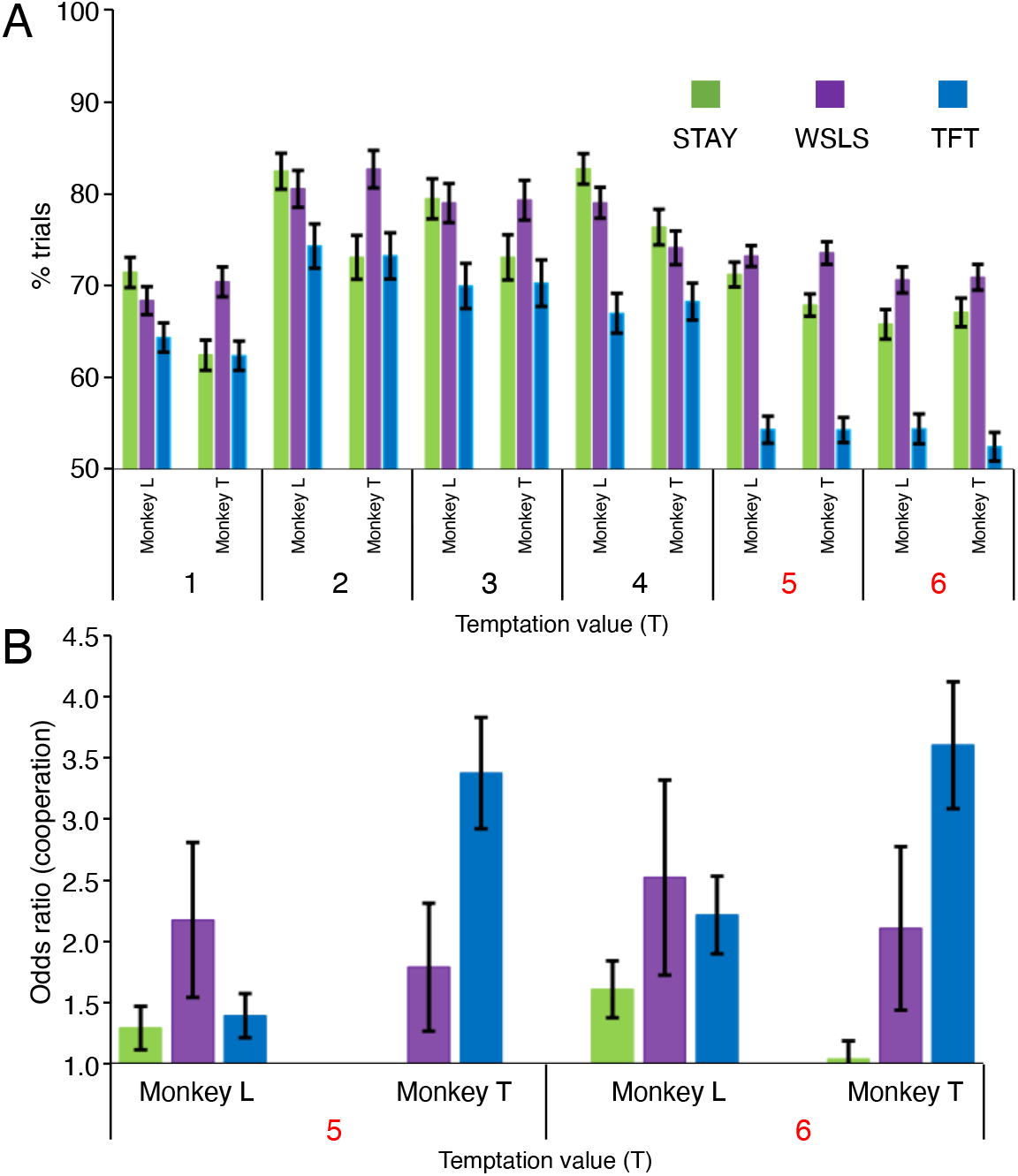
Strategies used in coordination and cooperation games: persistence (STAY), win-stay lose-shift (WSLS) and tit-for-tat (TFT). (*A*) Use of the different strategies by Monkeys L and T. Random choice would result in 50% selection of trials. Monkey L used STAY and WSLS more often than TFT, whereas Monkey T used mostly WSLS. Both animals maintained use of STAY and WSLS but decreased use of TFT substantially when passing from coordination games (Temptation = 1 – 4; black) to cooperation games (T = 5 and 6; red). Error bars show 95% confidence intervals. (*B*) Increase of cooperation with specific strategies. WSLS and TFT consistently increased cooperation more than STAY in the cooperation games.

## Discussion

We tested rhesus monkeys in a series of six games whose payoff matrices were identical except for the temptation payoff that increased across the games (Fig. 1A-D). This temptation payoff defined the transitioned from four coordination games to two cooperation games. The payoff matrix of formal cooperation games (PD) is characterized by the temptation payoff that exceeds all other payoffs and encourages defection that challenges cooperation (Fig. 1B). The animals showed strong performance in the coordination games that lead to highest payoff by common choice; their cooperation performance was somewhat lower but nevertheless substantial and stable despite the challenging higher defection payoff (Fig. 2A-D). Cooperation developed gradually over several weeks and months (Figs. 2E-F) but remained lower than performance in the coordination games (Fig. 3A, B). The degraded cooperation with reduced visual contact between the animals emphasized the social nature of the games (Fig. 3C). The animals’ choices seemed to follow common intuition; in the cooperation games, the first-choosing animal chose the commonly rewarded option less frequently than in the coordination games (Fig. 4A), which may reflect the low non-cooperative payoff in cooperation games. Further, choice of the cooperation option by the first player increased the chance of reciprocation by the second player (Fig. 4B, C). Correspondingly, successful cooperation in the current trials increased cooperation in the next several trials, suggesting that cooperation induced further cooperation and thus was behaviorally infectious (Fig. 4D). Reward accumulated more with mutual coordination compared to individually different choices, which confirms the beneficial nature of working together; more interestingly, reward payoff increased also with mutual cooperation despite the possibility of defection, thus demonstrating gain from engaging in pro-social rather than selfish behavior (fig 5). This gain may have been due to the acquisition of beneficial social strategies that increased cooperation (Fig. 6). Thus, rhesus monkeys showed solid and beneficial cooperation behavior in formal economic games following gradual, single-variable transition from coordination games.

In sum, the studies by others and ourselves demonstrated variable behavior in cooperation games, in contrast to the more stable choices in coordination games devoid of attractive defection. Cooperation in our experiments likely benefitted from extensive experience with coordination games that familiarized the animals with the benefit from choosing the cooperative option and allowed them to develop mutually beneficial strategies. However, the low number of monkeys in all these studies would prevent us from drawing strong conclusions.

### Methodological aspects

The single difference between our coordination and cooperation games consisted in the temptation payoff (T) for choosing the defection option (D) when the other monkey chose the cooperation option (C). By definition, the temptation payoff in a cooperation game exceeds all other payoffs (along with the payoff hierarchy shown in Fig. 1B), whereas the temptation payoff in a coordination game does not exceed any other payoff (Fig. 1C). Despite this higher temptation payoff, the animals succeeded to cooperate above chance in the cooperation games. Nevertheless, cooperation decreased with increasing temptation payoff between the coordination and the cooperation games, a difference that was larger between coordination and cooperative games than it was within these two game types (Fig. 3B). The drop in cooperation with increasing temptation payoff suggests that the animals detected the transitions to the more costly cooperation. However, the animals still maintained significant cooperation in the cooperation games (Fig. 3B), which may reflect familiarity with the lower temptation payoffs in the coordination games that elicited less defection. The good performance in the cooperation games may be a result of having experienced the coordination games with closely related payoff matrices that might have made the animals resilient to temptation. Alternatively, the coordination games might have primed the animals to focus on the cooperation option whose common choice led to the highest payoff, analogous to the direct priming with maximum reward for common choice (Stephens et al. 2002).

The ultimate gain in a cooperation game requires an initial loss. In our PD, unilateral defection by only one player paid always more reward (5 or 6 units) than unilateral cooperation (1 unit), whereas mutual defection (2 units) was less rewarding than mutual cooperation (4 units) (Fig. 1D). Thus, the initial higher reward with unilateral defection needs to be overcome by the experience of ultimately higher reward from mutual cooperation. The experience of the higher reward from cooperation would encourage a behavior associated with initially smaller payoff from unilateral choice. Thus, the challenge in cooperation, as represented in the formal PD, is to overcome the initial individually lower reward for the ultimately higher common gain.

While ‘cooperate’ choices in the coordination games were robust irrespective of the temptation payoff, ‘cooperate’ choices decreased in cooperation games with increasing temptation payoff. Thus, the temptation payoff in the cooperation games constituted a challenging defection option; the less the animals cooperated unilaterally, the more they were rewarded. Thus, by choosing the more rewarded defection option, the animals acted ‘as if’ they were maximizing reward; the animals ‘knew what they were doing’, and their choices seemed meaningful. Nevertheless, their substantial, above-chance cooperation against selfish higher-paying defection indicated their understanding of the long-term gain of cooperation.

### General game behavior

The obtained cooperation results confirm the notion that cooperation is beneficial in iterated PD. By contrast, the single-shot PD has its Nash equilibrium in defection (nobody gains by using a different strategy). The difference in optimal behavior is surprising, has been long debated, and seems quite controversial. One explanation is related to the length of the iteration. With predetermined finite sequences with short and well known length, an agent can iteratively backtrack from the last choice via the preceding few choices to the first choice and behave as if it were a single-shot PD, thus choosing defection to maximize own reward. However, in sequences with unknown and extensive lengths, backtracking to the first choice is more difficult (Dixit & Skeath 2004). This characteristic may be a reason why cooperation can be the prime characteristic of iterated PD. The beneficial cooperation seen in the current experiments confirm the validity of this notion in rhesus monkeys.

Our animals showed less frequent choice of the ‘cooperate’ option in the cooperation games compared to the coordination games. This observation corresponds to a previously described cooperation decline from two coordination games to the iterated PD, although the coordination games differed from those tested presently (Smith et al. 2019). The similar pattern of cooperation decline confirms empirically the intuitive notion that the the highest rewarded temptation payoff defining PD games constitutes the main challenge to cooperation.

The robust cooperation in our iterated PDs surpassed the degree of cooperation seen in a previous study that did not implement a transition from coordination to cooperation games (Haroush & Williams 2015). These authors reported 17% common cooperation choices, which is below the 28 – 68% cooperation choices seen presently (Fig. 3B). Despite the experimental differences of such complex social studies between laboratories, one factor for the presently observed more frequent cooperation may be our priming of cooperation by extensive experience with coordination games. In particular, the experience with the cooperation games may have increased the animals’ preference of the ‘cooperate’ option to its ‘defect’ alternative.

### Use of social information

Our experimental setup allowed the monkeys to make choices while being able to see their opponent and its choice in every trial before making their own choice. Seeing the opponent’s choices that affect the outcome of the own choice emphasizes the social nature of the task; it allows the animals to learn the contingencies of the game and its outcomes and appreciate the consequences of their own choices in dependence on the opponent’s choices. And importantly, seeing the influence of the opponent’s choices on their own outcome should prevent the animals from assuming simple stochasticity of the outcomes; they learn that the other animal has an influence on the own outcome. Without such information, for example when replacing animals by computer opponents or placing them in separate rooms, the animals might assume that the choices are stochastic and then use the probabilities of outcomes as a measure to determine their choices.

The visual interaction allowed each animal to respond to the choice of the opponent, thus minimising losses from unilateral cooperation. Indeed, a visual barrier between the two animals that prevented them to see the opponent’s options and choice resulted in rapid decline of cooperation (wall control; Fig. 3C). These results correspond to the previously reported drop in individual choice of the cooperation option when replacing the opponent monkey by a computer (from 35%% to 19%; Haroush & Williams 2015). A similar drop of individual choice of the cooperation option was observed when placing the opponent monkey in another room (from 35% to 14%; Haroush & Williams 2015). The replacement of a biological partner by a computer opponent and the animals’ placement in separate rooms reduce the social aspect more than the simple blocking of the view of each other’s options and choices achieved by our wall control. Nevertheless, irrespective of the degree of reduction of social interaction, the results demonstrate the importance of social aspects in these formal, and somewhat abstract and reductionist, economic tasks.

### Strategies

Our monkeys cooperated even when they could see each other’s options and choice, which might have made them more vulnerable to exploitation and defection. Despite frequent defection in the face of the opponent’s cooperation, the animals cooperated in most trials: when the first moving animal chose to cooperate, the other animal often chose also to cooperate. This result held for all animals and all games (Figs. 2, 4).

Formal analyses of our animals’ choices revealed use of three major strategies (Fig. 6): persistence (STAY), win-stay lose-shift (WSLS) and tit-for-tat (TFT). STAY consists of choosing the same option as before irrespective of the outcome, whereas WSLS consists of choosing the same option after receiving a good outcome but choosing the alternative after a lesser outcome. Thus, players using STAY perseverate without taking the other player’s choice into account, and players using WSLS respond to the outcome of their own choice that also depends on the other’s choice. Thus, the two strategies are either not social at all (STAY) or social only to a limited extent (WSLS).

By contrast, TFT consists of reciprocation and thus constitutes a fully social strategy. The player replicates the opponent’s play, cooperating after a cooperation until the other player defects, and defecting after a defection until the other player cooperates. As mutual defection results in lower outcome than mutual cooperation, players should aim for mutual cooperation; they should occasionally even offer to cooperate in the face of the other player’s defection. This behavior may result in long periods of being unilaterally defected that is beneficial for the opponent but provides suboptimal own outcome (see payoff matrix in Fig. 1D) that is only justified by the later gain from mutual cooperation. In contrast to TFT, defection in WSLS would lead after only one trial to stable cooperation. Thus, recovery of mutual cooperation after defection is more costly in TFT than WSLS, which may be a reason why WSLS outperforms TFT in humans (Nowak & Sigmund 1993).

These strategies are also found in our monkeys (Fig. 6A). First, the animals used STAY and WSLS rather consistently in all games, whereas they used TFT much more rarely in both cooperation games. Indeed, WSLS resulted in more cooperation than the other strategies (Fig. 6B). An earlier study had shown substantial variability among individual monkeys to establish stable strategies, but the animals expressing stable strategies did show reciprocal cooperation and defection compatible with TFT (Smith et al. 2019). In another study, monkeys tended to continue cooperating when the opponent had cooperated but tended to defect less when the opponent had defected, suggesting some correspondence to both WSLS and TFT that significantly exceeded random behavior (Haroush & Williams 2015). Their reciprocation depended on the social situation by decreasing when one player was replaced by a computer or when performing in different rooms.

## Acknowledgements

We thank Aled David and Christina Thompson for animal and technical support and Colin Camerer, Raymundo Báez-Mendoza, Alexandre Pastor-Bernier and Fabian Grabenhorst for discussions on game theory and the design of this experiment. This study was supported by Wellcome Trust (WT 095495, WT 204811, WT 206207), European Research Council (ERC; 293549) and US National Institutes of Mental Health (NIMH) Caltech Conte Center (P50MH094258). For the purpose of Open Access, the authors have applied a CC BY public copyright licence to any Author Accepted Manuscript version arising from this submission.

